# The NanoZoomer Connectomics Pipeline for Tracer Injection Studies of the Marmoset Brain

**DOI:** 10.1101/748376

**Authors:** Alexander Woodward, Rui Gong, Hiroshi Abe, Ken Nakae, Junichi Hata, Henrik Skibbe, Yoko Yamaguchi, Shin Ishii, Hideyuki Okano, Tetsuo Yamamori, Noritaka Ichinohe

## Abstract

We describe our connectomics pipeline for processing tracer injection data for the brain of the common marmoset (*Callithrix jacchus*). Brain sections were imaged using a batch slide scanner (NanoZoomer 2.0-HT) and we used artificial intelligence to precisely segment the anterograde tracer signal from the background in the fluorescence images. The shape of each brain was reconstructed by reference to a block-face and all data was mapped into a common 3D brain space with atlas and 2D cortical flat map. To overcome the effect of using a single template atlas to specify cortical boundaries, each brain was cytoarchitectonically annotated and used for making an individual 3D atlas. Registration between the individual and common brain cortical boundaries in the flat map space was done to absorb the variation of each brain and precisely map all tracer injection data into one cortical brain space. We describe the methodology of our pipeline and analyze tracer segmentation and brain registration accuracy. Results show our pipeline can successfully process and normalize tracer injection experiments into a common space, making it suitable for large-scale connectomics studies with a focus on the cerebral cortex.

## 1 Introduction

Across the world, large-scale brain mapping projects are becoming increasingly common (Markram et al. (2011); Jorgenson et al. (2015); Okano et al. (2016); Grillner et al. (2016); Poo et al. (2016); Lin et al. (2019)). Successfully carrying out these projects requires an integrated and inter-disciplinary approach that leverages the scalability and automation afforded by computational pipelines. To map the brain of the common marmoset (*Callithrix jacchus*) we describe our connectomics pipeline for tracer injection studies using a NanoZoomer 2.0-HT batch slide-scanner.

Our novelty comes firstly from artificial intelligence being used for the first time on NanoZoomer fluorescence images of the common marmoset brain to precisely segment anterograde tracer signal. This approach overcomes low-frequency intensity distortions and bright non-tracer features that precluded the use of standard thresholding. Secondly, to overcome bias from using a single brain template to specify cortical boundaries we created an individual atlas for each brain. Cytoarchitectonic annotations by a neuroanatomist were automatically processed and converted into 3D. All brain image data was non-linearly mapped into a new common 3D brain space based on a population average magnetic resonance imaging (MRI) brain mapped with a 3D marmoset brain atlas. The cortex of the individual brain was then projected into a 2D cortical flat map space associated with the common 3D brain space. Finally, 2D registration between the individual and common brain cortical boundaries was done to absorb individual brain variation and precisely map all tracer injection data into one common cortical brain space.

The basic steps involved in tracer injection connectomics are as follows: raw data of the brain is reconstructed into 3D, the injection site and tracer signal are identified, and this data is mapped in 3D with a brain atlas. Then a connectivity matrix describing the structural connectivity between brain regions can be calculated. For the common marmoset brain this fiber connectivity is still poorly understood and such an approach can help provide a foundation for developing a neurobiological understanding of higher brain function.

In the studies used for this pipeline, viral anterograde tracers were used to infect neuronal cells, causing them and their axon fibers to fluoresce when imaged using fluorescence microscopy. We build upon the results described in Abe et al. (2017) which uses a method for the 3D reconstruction of marmoset brain section images by registering to corresponding block-face images. By computationally reconstructing the brain sections into 3D we can discover the fiber projection patterns between different anatomical regions.

We intend to integrate our results with other tracer injection based connectomics pipelines through our shared common brain space, contributing to a future marmoset whole-brain connectome. This includes tracer injection studies using the TissueCyte serial two-photon tomography system (Skibbe et al. (2019)), in which the approach for using artificial intelligence for tracer segmentation was established. Thus, we call our pipeline the NanoZoomer Connectomics Pipeline.

## 2 Materials and methods

Figure 1 describes the pipeline workflow. Experimental data for each brain (described in Section 2.1) was acquired and transferred across the internet to be stored on a central server with large-scale shared storage (LSSS) of 1.2 PB using the general parallel file system (GPFS) file system. The server has a high-performance cluster (HPC) system that we used to process the experimental data in a batch manner. The HPC system has 18 computing nodes (16 × Dell PowerEdge M620 and 2 × Dell PowerEdge R720), each node having two Intel Xeon E5-2680v2 (2.8GHz, 10 Cores) CPUs and 256 GB of memory, giving a total of 360 CPU cores. The HPC runs the Linux Rocks Cluster 6.2 OS (CentOS 6.9 x86-64). The HPC login node was a Dell PowerEdge R620 (Intel Xeon E5-2630v2 (2.6GHz, 6 Cores) × 2 with 64 GB of RAM). This was used for job scheduling using the Open Grid Scheduler/Grid Engine 2011.11p1 (OGS/GE) (Sun Grid Engine(SGE) based) job scheduler. The artificial intelligence pipeline parts were trained and processed on a Dell Precision 7920 desktop machine with two Nvidia GPUs (RTX 2080 Ti and GTX 1080 Ti), running Ubuntu OS 16.04 and connected to the server’s LSSS via 10 Gb Ethernet.

**Fig. 1.**
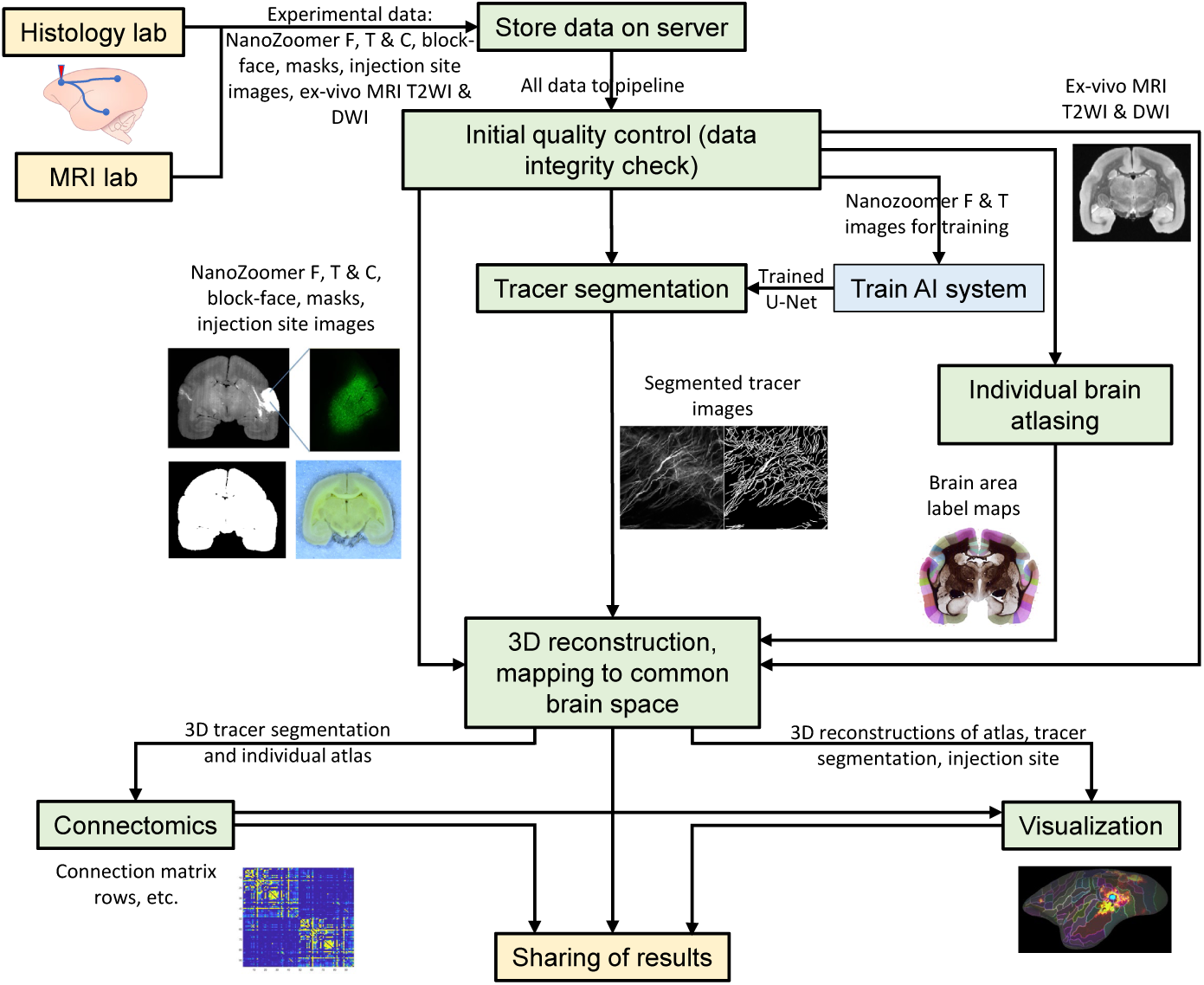
The pipeline workflow where each green box represents a process. Experimental data is described in Section 2.1, tracer segmentation in Section 2.3, individual brain atlasing in Section 2.4, 3D reconstruction and mapping to the common brain space in Section 2.5, examples of connectomics and visualization are shown in Section 3.

An automatic quality control step was used to check the integrity of received experimental data, making sure that all of the required images and meta-data existed for processing at later stages. This check either returned a pass or fail. Visual checks were also done on the pipeline output to see if any adjustments or corrections needed to be made. These were communicated to the experimental labs and the data was modified and re-uploaded as appropriate. Next the data processing begins: the tracer signal was segmented (Section 2.3) and individual brain region delineations were processed into 2D label maps (Section 2.4. Then data was reconstruction into 3D, namely the block-face (Section 2.5.1, which is used to map with the common brain space, the fluorescence images (Section 2.5.2, the Myelin and label map images (Section 2.5.3), and the injection site (Section 2.5.4). By mapping data into a common brain space we have the opportunity to integrate connectomics information from a wide number of sources. Furthermore, because there is a transformation path from the original raw data sets all the way to the common space we can map data from it into the space of any of the individual brains. Finally, the processed data can be used to calculate connectomics information from each injection.

Our pipeline was developed using only open-source tools and libraries. The core was written in Python with the Nipype framework (Gorgolewski et al. (2011)) managing parallel processing jobs. Nipype made it straightforward to construct a complex pipeline and Python has some optimized image processing and machine learning algorithms suitable for neuroinformatics tasks. For 2D image processing we used the Python bindings for the OpenCV (Bradski (2000)) and SimpleITK (Lowekamp et al. (2013)) libraries. For 3D image processing we used SimpleITK and the ANTs Normalization Tools (Avants et al. (2011)). For all 2D and 3D image registration tasks ANTs was used. For non-linear registration we used the ANTs implementation of the SyN algorithm (Avants et al. (2008)), a top performer for medical image registration (Klein et al. (2009)). The artificial intelligence parts used Keras (Chollet et al. (2015)) with a Tensorflow backend (Abadi et al. (2015)). Configuration settings for the pipeline were stored in JSON files or simple text files. All of the generated image data was saved in 2D TIFF or 3D NIfTI formats, making it easy to export results into a wide number of existing neuroscience tools. Our results are also compatible with the Human Connectome Project’s Connectome Workbench software (Marcus et al., 2011).

### 2.1 Experimental data

#### 2.1.1 Brain sectioning and block-face imaging

Dry ice was used to freeze the brain and a sliding microtome (SM2010R, Leica) was used to slice the coronal brain section at 50*µm* thickness (z-sampling interval). After a section was sliced an image of the brain block-face was taken using a camera (D5500, Nikon Corporation, Tokyo, Japan) with a macro lens (AF-S NIKKOR 1835 mm f/3.54.5G ED, Nikon, Tokyo, Japan) and fixed to a copy stand. This produced around 600 brain sections on average. Since the camera was fixed during acquisition, the block-face images can be computationally stacked to recover the 3D shape of the brain.

Brain boundaries in the block-face images were often faint due to the similarity in pixel value with the dry ice surrounding it. Therefore the brain boundary was manually delineated and binary mask images were created for every fourth section were created. Intermediate masks were created by solving Laplace’s equation with the nearest adjacent masks acting as boundary conditions. We plan to look at using color histogram segmentation to remove the background region automatically.

Brain sections were sequentially grouped into sets of three, where the first section was used for Myelin staining, the second for Nissl staining and the third for fluorescence imaging. Thus there is a one-to-one correspondence between a block-face image and a Myelin, Nissl, or fluorescence image that can be used for recovering the brain shape.

#### 2.1.2 NanoZoomer 2.0-HT fluorescence images

Two anterograde tracers which express green and red fluorescent proteins and have maximal fluorescence at different frequencies of light were injected into the left hemisphere of each brain. The viral tracer was a mixture of AAV1-Thy1S-tTA (titer: 1×109 vector genomes (vg/*µl*)) and AAV1-TRE-hrGFP (5×109 vg/*µl*). A batch process slide scanner (NanoZoomer 2.0-HT, Hamamatsu Photonics K.K., Hamamatsu, Japan; 20x objective, 455 nm/pixel) was used to capture the fluorescence images. Brain sections were scanned using FITC (green, channel F), TRITC (red, channel T), and an additional Cy5 (far red, channel C) filter (LED-DA/FI/TR/Cy5-4X-A-OMF, Semrock, Inc, NY). The C channel was used to capture the auto-fluorescence of the brain tissue. For each channel an average of 200 images were generated and saved as 16-bit TIFF images. A copy of the fluorescence images were reduced to the block-face image resolution and binary image masks were manually created to mask the area outside of the brain region of interest.

#### 2.1.3 Histological staining and 2D cortical region delineation

Myelin and Nissl staining was used for cytoarchitectonic delineation of brain regions. The 2D boundaries of cortical regions were drawn over each Myelin image and labels of the anatomical abbreviations for each region were added (following the nomenclature of Paxinos et al. 2012). Masks for the brain section boundary in each Myelin image were created by non-linear registration of the already created block-face masks into the Myelin image space. To reduce manual labor the gray/white matter boundaries were drawn for a sub-set of the Myelin images and mapped to adjacent images using non-linear image registration. Drawing was carried out on a graphics tablet (Cintiq DTH-2200/K1, Wacom, Saitama, Japan) and image editing software (CorelDRAW essentials X6, Ottawa, Canada). Region delineation data was saved in the SVG vector file format.

#### 2.1.4 Injection site images

To capture fine axonal projection patterns the sensitivity of the NanoZoomer scanner was set to a high level that unfortunately caused saturation at the injection site (the region of infected cell bodies that show strong fluorescence when imaged). This made it difficult to precisely estimate the injection site volume from the NanoZoomer scanned images. To solve this issue, images of the injection site region for the F and T channels were taken using an epifluoresence microcrope (BZ-X700, Keyence, Osaka, Japan) with a 4x objective (CFI Plan Apo lamda, NIKON corporation, Tokyo, Japan). The spatial resolution of each image was recorded using the camera viewer software. Masks of the tracer injection site were manually created for each image showing bright cell bodies. We intend to use this data for train an automated artificial intelligence approach for injection site detection.

#### 2.1.5 Ex-vivo MRI T2-weighted images and diffusion-weighted images

A Bruker BioSpec 9.4T MRI machine (Biospin GmbH, Ettlingen, Germany) and a solenoid coil with an 28-mm inner diameter was used to acquire exvivo MRI T2-weighted images (T2WI) and ex-vivo diffusion-weighted imaging (DWI) data for each marmoset brain. For T2WI, a rapid acquisition with relaxation enhancement sequence was used with the following parameters: TR = 10000 ms, TE = 29.36 ms, flip angle = 90^°^, resolution = 100 × 100 × 200*µm*^3^, NA = 16, and scan time = 3 hour 20 min. For DWI, spin-echo type multi shot echo planar imaging was used for the assessment of diffusion properties with the following parameters: TR = 4000 ms, TE = 28.39 ms, resolution = 200 × 200 × 200*µm*^3^, *δ* = 7 ms, *Δ* = 14 ms, b-value = 1000, 3000, 5000 s/mm^2^ in 128 diffusion directions (plus 6 b0 images), k-space segmentation = 10, NA = 2, and scan time = 8 hour 37 min.

MRI T2WI was chosen as a common modality for image registration of exvivo brain experiments and DWI can be used for comparing tracer injection results with DWI based tractography. A population based ex-vivo MRI T2WI brain was generated from the individual MRI scans and was used for the common brain space in this work.

### 2.2 Common brain space using a population average MRI T2WI and a 3D digital marmoset atlas

The common brain space was composed of a population average ex-vivo MRI T2WI contrast mapped with the freely downloadable Brain/MINDS 3D digital marmoset brain atlas (Woodward et al. (2018a,b)). The population average MRI was constructed from 25 individual brain scans by iteratively applying linear and non-linear registration and averaging the transformation files until convergence. Data was resampled before processing to give an average brain space with an isotropic spatial resolution of 100 ×100 × 100*µm*^3^. The registration procedure gave a brain shape with a high signal-to-noise ratio compared to an individual MRI scan. The average MRI was then AC-PC aligned within an RAS (Right-Anterior-Superior) coordinate system.

The Brain/MINDS atlas is a multi-modal atlas with individual MRI T2WI and co-registered Nissl histology data Woodward et al. (2018a). Anatomical regions were delineated in original high-resolution Nissl image data which was then computationally mapped into the 3D shape of the MRI T2WI. The atlas has 280 brain region delineations per hemisphere, giving 560 regions in total. For each hemisphere there are 114 cortical regions grouped into 14 cortical areas (isocortex), with an additional 4 non-isocortex regions. In addition to the cortical region segmentation, gray, white, and mid-cortical segmentations were defined in the atlas. We precisely mapped the Brain/MINDS 3D digital marmoset atlas to the population average MRI brain using linear and nonlinear registration by using both the MRI contrasts and manually defined cortical and gray matter segmentations. We used the mid-cortical segmentation mapped into the population average space to create a 2D cortical flat map space. This was created using the Caret software (Van Essen et al. (2001)) and is compatible with the Connectome Workbench software (Marcus et al. (2011)). We can map any data from the pipeline into the cortical flat map space for intuitive 2D viewing. Fig. 2 visualizes the common brain space with cortical flat map.

**Fig. 2.**
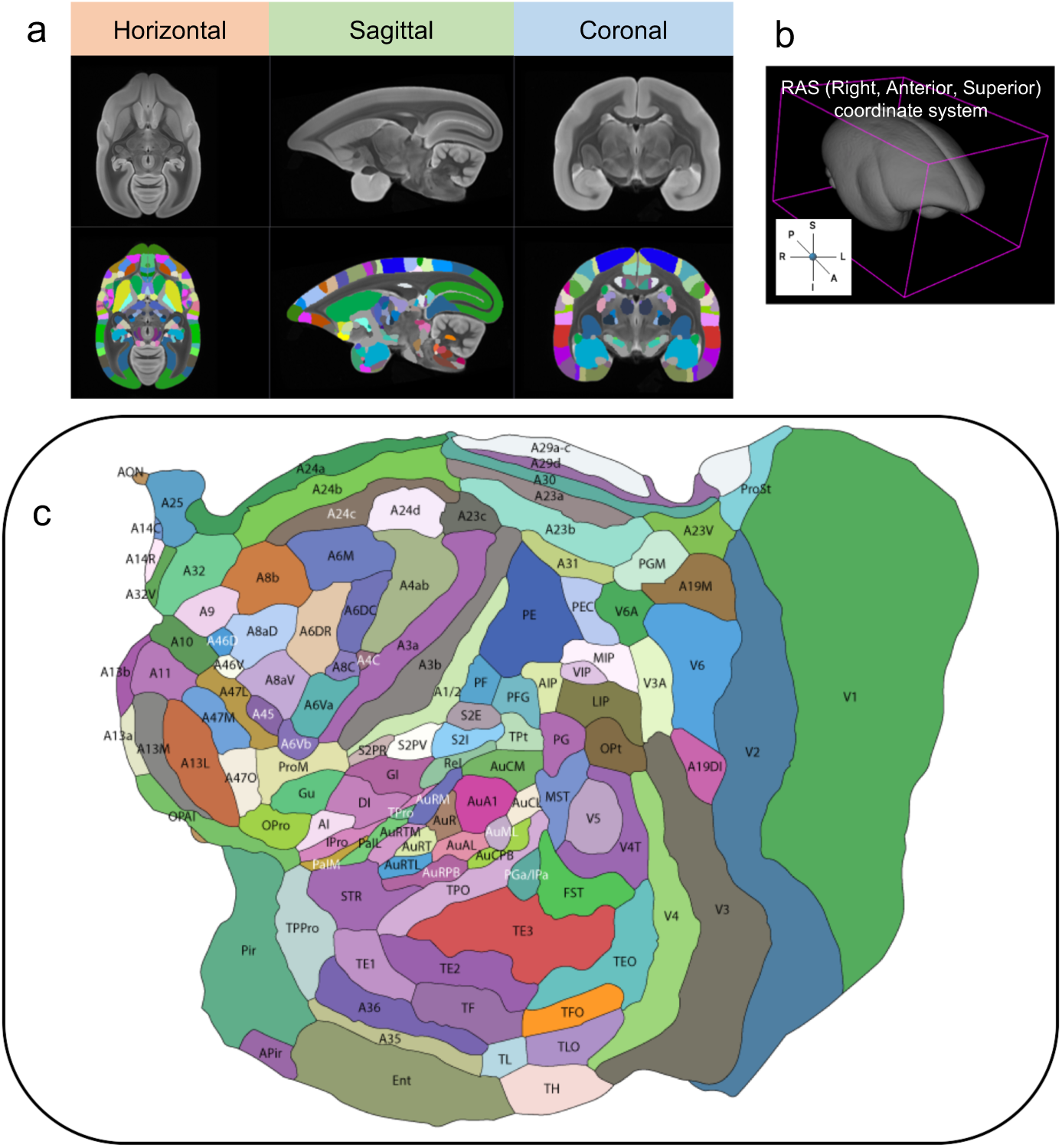
The common brain space consisting of the Brain/MINDS 3D digital marmoset atlas mapped to a population average ex-vivo MRI T2WI brain. The left hemisphere cortical flat map for the Brain/MINDS 3D digital marmoset atlas.

### 2.3 Tracer signal segmentation using the U-Net deep network architecture

To identify the fiber projection pathways of injected tracer and to measure the amount of tracer in each region we needed a robust method to process the tracer images. Non-tracer signal in the images from the NanoZoomer 2.0-HT batch slide scanner included image artifacts, infected cell bodies, blood vessels, or non-linear variation in the intensity of brain tissue regions. Traditional methods such as intensity thresholding can fail to separate the background from tracer due to the non-linear illumination or to identify very weak tracer signals.

We used the U-Net deep learning architecture (Ronneberger et al. (2015)) to try to overcome the challenges of segmenting tracer signals from brain images with non-linear illumination. This is a popular convolutional neural network architecture used mainly for biomedical image segmentation problems where the number of training images can be limited. We used a freely available Tensorflow/Keras implementation described in Taashi-s (2018).

A trained U-Net takes as input the images that should be segmented and outputs a probability map that can be used to classify all pixels in an image into background or tracer. For training data, we chose fluorescence images from several brain slices along with manually created masks for each image, with a value of 0 for the background and 1 for the tracer signal.

To create training data we selected a sub-set of images from the T channel, each with a resolution about 55000 × 50000 pixels, and cropped them into 1000 × 1000 pixel sub-images. From all of the sub-images, we selected and manually drew segmentation masks for 130 sub-images, and used another 163 background images where there was no tracer signal. We then sliced each of the sub-images into four 500 × 500 pixels image before input into the neural network for training.

For data augmentation, we applied rotations and gamma augmentation between 0.5 and 1.5. This was necessary due to limited training data, and a large training data set is necessary for good performance in a neural network.

Choosing good hyperparameters is an essential part of neural network training as they control the behavior and performance of the network. The hyperparameters used for training were: 5 encode and decode layers, 32 feature channels at the top layer, 512 feature channels at the bottom layer, with padding at each layer to ensure even-sized feature maps. We used the ‘Adam’ optimizer (Kingma and Ba (2014)) with a learning rate of 0.00001. The values of each input data were first standardized with a zero mean and standard deviation of one before beginning the training.

To prevent overfitting to the training data, where the network tries to memorize the input data rather than learn features from the input data, the network applied both batch normalization (Ioffe and Szegedy (2015)) and dropout layers (Srivastava et al. (2014)) with a dropout rate of 0.6.

Through data augmentation a total of 4688 images were used to train the neural network. The training data was split into 80% training and 20% validation. The validation set was a subset of images that were not used during the training of the network. The purpose of the validation set was to monitor the prediction accuracy of the training. For example, if the loss function from training was decreasing and the loss function from validation was not decreasing or sometimes increasing, it told us the network was overfitting the training data. We used binary cross-entropy as the loss function to monitor the error in training and prediction, since we expecting binary classification from our network.

Our training was done on a multi-GPU model using an Nvidia RTX 2080 Ti and an Nvidia GTX 1080 Ti graphics card. The network was trained for 500 epochs, taking just under 42 hours. An example of our trained U-Net applied to a previously unseen image is shown in Fig. 3. Impressively, the network managed to segment the tracer signal from the background while dealing with non-linear illumination and different image artifacts. It was also able to capture faint tracer signals that can barely be seen by eye. This can be seen in the visually denser amount of segmented tracer signal compared to what is apparent in the original image, especially in the bottom-right region of the image where a large amount of weak but dense tracer signal existed.

**Fig. 3.**
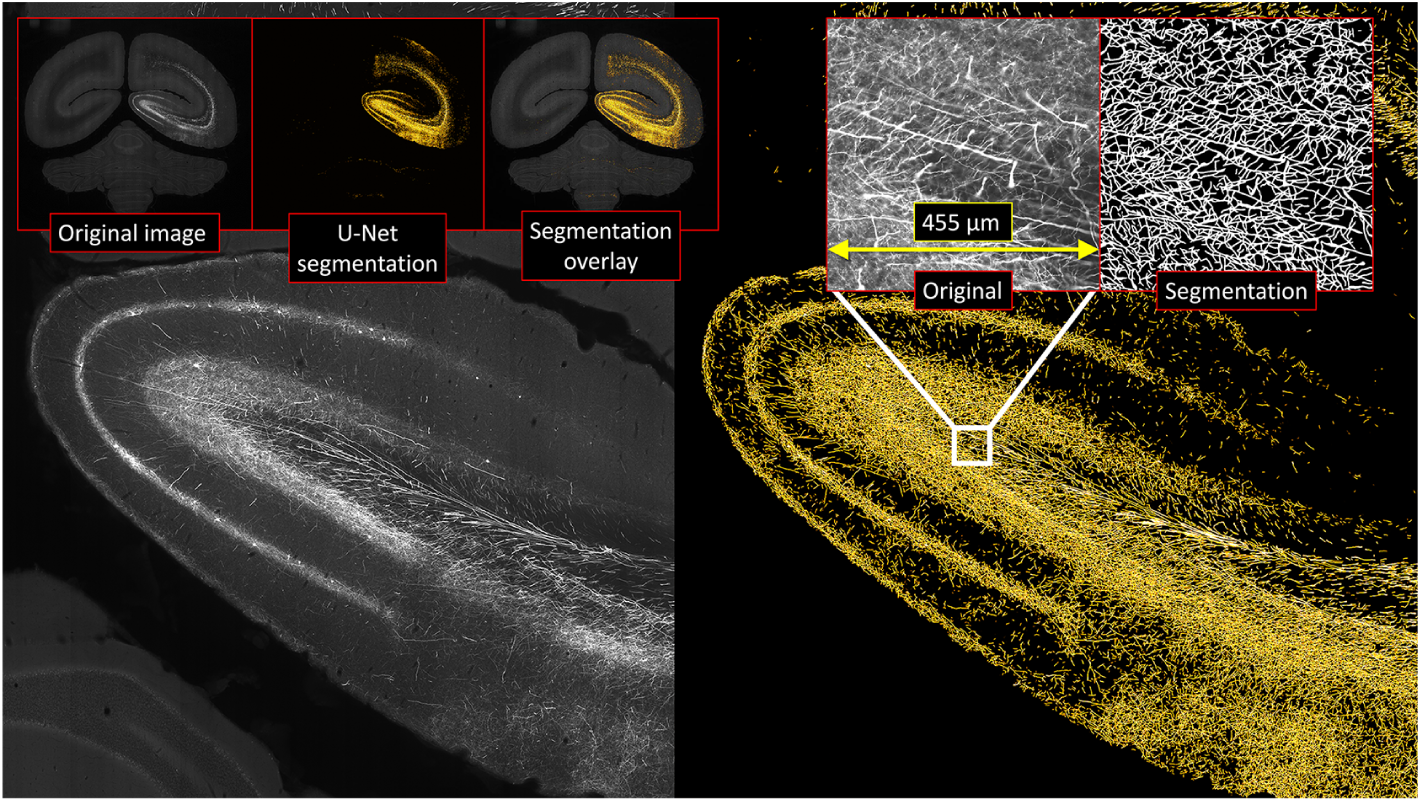
An example tracer segmentation result on a previously unseen image using our U-Net approach, showing a close-up view of a coronal slice of the left hemisphere occipital lobe. On the left is the brain auto-fluorescence and on the right is the segmented tracer signal result. The inset images above the segmentation on the right show a close up of the tracer signal, before and after segmentation, for a small sub-region.

### 2.4 Conversion of 2D cortical delineations into label maps

The boundaries of delineated 2D cortical regions had to be filled with a brain region ID before 3D reconstruction. The SVG files were first rasterized and the 2D positions of region labels were used as seed points to run a flood fill algorithm in each anatomical region. This process generates 2D label maps, where the pixels of each brain region had a value corresponding to a brain region ID in the Brain/MINDS atlas specification. Flood-filling the regions required that boundaries were entirely closed and the SVG processing procedure modified cortical boundaries when needed so that they terminated at the brain or gray/white matter boundaries.

### 2.5 3D reconstruction of data and mapping to the common brain space

#### 2.5.1 Block-face 3D reconstruction and mapping to common brain space

Block-face images were resized to 500 × 500 pixels and then converted to grayscale. The operating resolution was chosen as a compromise for carrying out subsequent 3D image registration in a reasonable amount of time. The block-face images were masked and computationally stacked into a 3D volume. The individual brain shape was easily recovered since all block-face images were aligned. This shape was used as a reference for aligning the slide-scanned fluorescence images and Myelin stained images.

We used linear and non-linear image registration of the population average MRI T2WI to the 3D reconstructed block-face, establishing a mapping between the two spaces. Repeating this procedure for each brain allowed us to integrate data across many tracer experiments. We also mapped the population average MRI T2WI with the individual MRI T2WI and DWI spaces for future analysis.

After registering the average MRI to the block-face we sliced it into a set of 2D images with a one-to-one correspondence with the block-face images. These were used for 2D registration with the fluorescence images. This was due to the higher contrast available in the population average MRI over the block-face.

#### 2.5.2 3D reconstruction of the NanoZoomer fluorescence contrast

The raw F, T, and C fluorescence channel images arrived in various sizes, having been cropped and saved using the NanoZoomer’s custom software. Since these images were extremely large (often between 1-3GB and up to around 50000 × 50000 pixels) they were first reduced to a size appropriate for doing 2D and 3D image registration in an expedient amount of time. A reduction was made to about 1% of the original resolution. This low resolution was chosen to roughly match the block-face image resolution. For analysis purposes the raw images were also transformed to 20% and 5% original resolution copies.

To standardize the image size for assembly into a 3D volume we created a 600 × 600 canvas and positioned each image and its corresponding mask into this space based on their image centers. The mask images were used to remove all image content outside of the brain region of interest. This was done for the F, T, and C channels.

We found that the F channel data was the most stable for driving image registration so it was used as the reference channel for 2D linear and non-linear image registration to the block-face registered average MRI images. The output of the 2D-to-2D registration generated 2D affine transformations and vector valued 2D images (warps) that described the linear and non-linear parts of the mapping respectively. This registration did not consider the alignment of brain structures across slices. We therefore implemented inter-slice consistency processing based around Gauss-Seidel iteration (Gaffling et al. (2015)). Here, adjacent images were warped to one another using non-linear image registration and their shapes were iteratively updated based on the calculated deformations. The procedure removed high-frequency distortions between slices while at the same time preserving the low-frequency shape. All transformation files for the aforemention procedures were saved and applied to the remaining T and C channels.

Finally, the transformed F, T and C channel image stacks were converted into NIfTI volumes. Then using the transformation established between the population average MRI and the block-face space (Section 2.5.1) we can map all fluorescence data into the common brain space.

#### 2.5.3 3D reconstruction of Myelin contrast and label map

The same approach for 3D reconstruction of the fluorescence images was used for the Myelin staining images. Since cortical region delineations were drawn over the Myelin images the same transformations were applied to reconstruct the label map images into 3D and create an individual atlas.

#### 2.5.4 Injection site 3D reconstruction

To deal with saturation at the injection site in the NanoZoomer images we processed images acquired from the BZ-X700 camera with a lower exposure setting where infected cell bodies were visible. Only the subset of images showing the injection site were captured and the indices of these images from the full set were recorded so that correspondences between NanoZoomer and BZ-X700 images was possible. This was carried out for the F and T channels. Affine 2D-to-2D image registration was performed between adjacent images and a reference image was chosen (the first image in the sequence). All images were transformed in 2D to align with the reference image by concatenating the transforms between the index of the current slice and the reference slice. By using the inter-slice step size (50*µm*) the aligned images could then be stacked into a 3D volume of the brain’s injection site sub-region.

For both the F and T channels the rough injection site region was located in the Nanozoomer 3D reconstruction through thresholding at a high value. The centroid of this region was calculated and used for placing the BZ-X700 injection region sub-volume. Since the naming of the injection site images corresponded with the NanoZoomer images we could precisely place the injection sub-region volume along the anterior-superior axis. Afterward, we visually checked the placement of the injection site and when necessary, manually adjusted the sub-volume’s position in-plane so that it lined up with surrounding features in the tissue fluorescence. The transformation pathway from 2D into the 3D common brain space was applied to the binary image masks delineating the injection site region in the BZ-X700 images. We were then able to recover a good estimate of the injection volume and its location in 3D.

#### 2.5.5 Alignment of individual atlas and common brain atlas in the cortical flat map space

The mapping procedure between individual and common brain space used block-face and MRI T2WI contrasts. This allowed us to normalize the brain shape so that the cortex boundary, white matter boundary and sub-cortical structures clearly aligned. However, variation of cortical boundaries between the mapped individual brain and the Brain/MINDS atlas mapped to the population average MRI still existed. To further absorb this variation we aligned cortical boundaries between the individual brain atlas and the common brain atlas in the 2D flat map space. We used non-linear registration to warp the 2D flat map space of the individual to the common space cortical boundaries. This projection also allows us to intuitively visualize cortical projection patterns in 2D. It remains future work to take the 2D flat map space alignment and use it to modify the 3D alignment in the common brain space.

## 3 Results

Before putting our pipeline into operation we trained our U-Net for tracer segmentation and constructed the common brain space using a population average MRI T2WI mapped with the Brain/MINDS 3D marmoset brain atlas. We then used our pipeline to completely process three brains (Cases 1, 2, and 3) and show the results in this section. We also assessed the accuracy of the 3D mapping of individual brains into the common space, and the U-Net tracer segmentation.

Figure 4 shows a 3D reconstruction of the atlas of an individual brain using the procedure described in Section 2.5.3. Cortical boundaries drawn in 2D were turned into label maps. The shape of the brain was reconstructed using the Myelin contrast mapped to the block-face allowing us to then turn the 2D label maps into 3D brain regions. Figure 5 shows the effectiveness of the 3D reconstruction procedure for fluorescence data (Case 2). Figure 5a shows a coronal view of the injection site in the NanoZoomer (saturated region) and BZ-X700 images (cell bodies are visible). Fig. 5b shows an example 3D reconstruction: the top row shows horizontal, sagittal and coronal views of stacking fluorescence sections (F channel) by their centroid. The bottom row shows the corresponding views after the sections have been registered to the block-face shape. The brain shape was successfully recovered from a set of initially unaligned brain section images. Finally, Fig. 5c shows horizontal, sagittal, coronal and 3D views of the injection site and segmented tracer signal mapped into the 3D common brain space. Fig. 6 shows results of the output of our pipeline for three brains. The segmented tracer, the individual atlas, and the injection site were mapped into the common brain space and then the cortical flat map space. The cortical boundaries between the individual and the common brain space were aligned in 2D. Data was visualized on the left cortical flat map and 3D mid-cortical surface. After 3D reconstruction and mapping of data we can progressively construct a brain connectivity matrix by quantifying tracer signal projections from each injection site (source) into different anatomical regions (target sites). It remains future work to clarify the best procedure to integrate and normalize different injection sites into a single connection matrix. However, we conducted preliminary connectomics calculations of cortical connectivity as a proof of concept. We defined the projection signal strength between a source and target region pair as *C*_*D*_ = *C*_*T*_ */C*_*R*_. Where *C*_*T*_ is the sum of tracer signal voxels that have non-zero value in the target region, and *C*_*R*_ is the sum of voxels of the injection area within the source region. *C*_*R*_ is used as a normalizing factor when combining data from multiple injections. An example connectivity result for Case 2 with an injection into region V5(MT) (T channel) is shown in Fig. 7. The regions that had connectivity were colored using a lookup table. Values were threshold to show only strongly connected regions where at least 10% of the region was filled with tracer signal. These values were then re-scaled for visualization purposes.

**Fig. 4.**
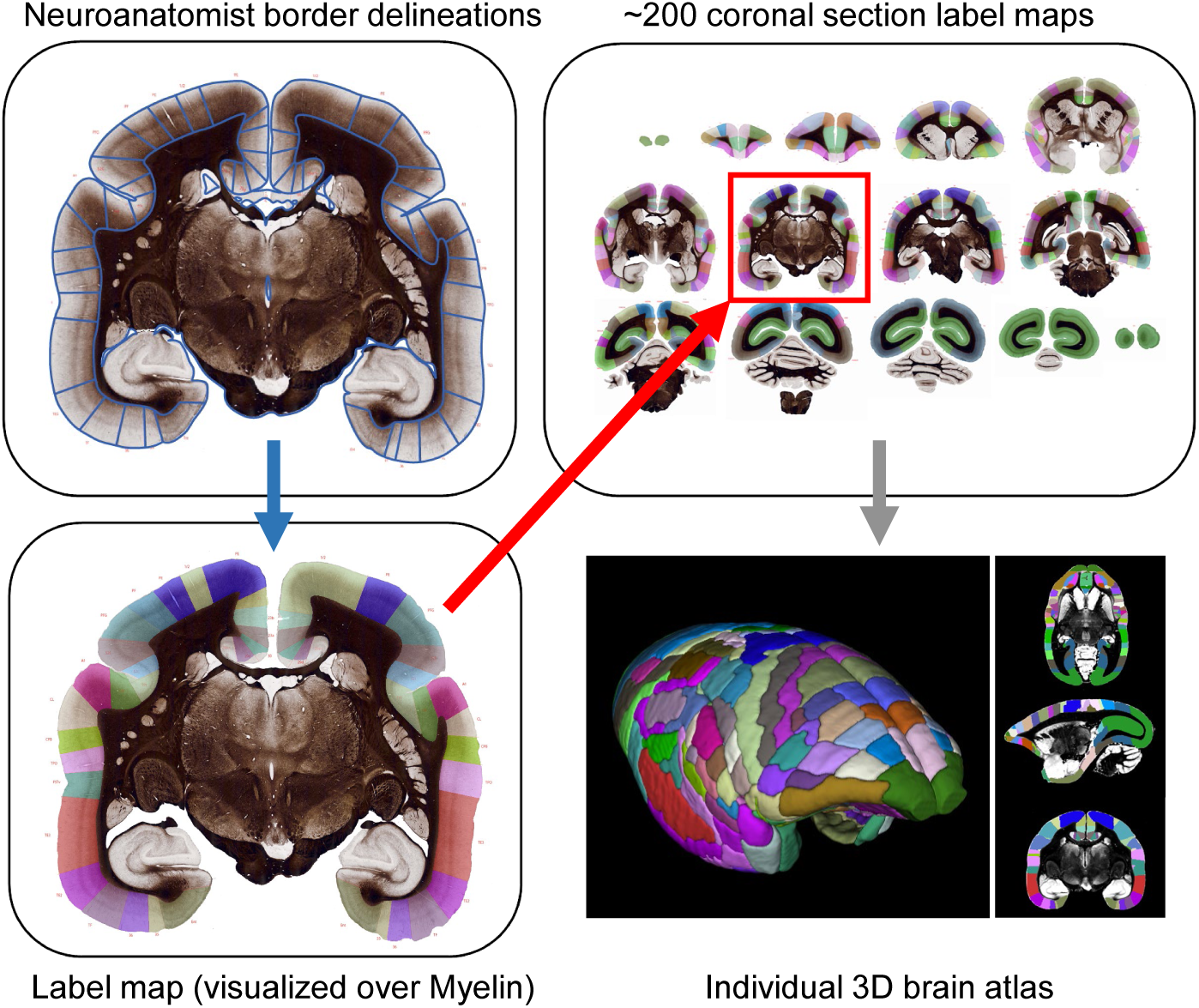
An example of the 3D reconstruction of an individual atlas. A slice showing the region delineations is given as input and this automatically converted into a raster label map. The set of all 2D label maps is processed and reconstructed into 3D.

**Fig. 5.**
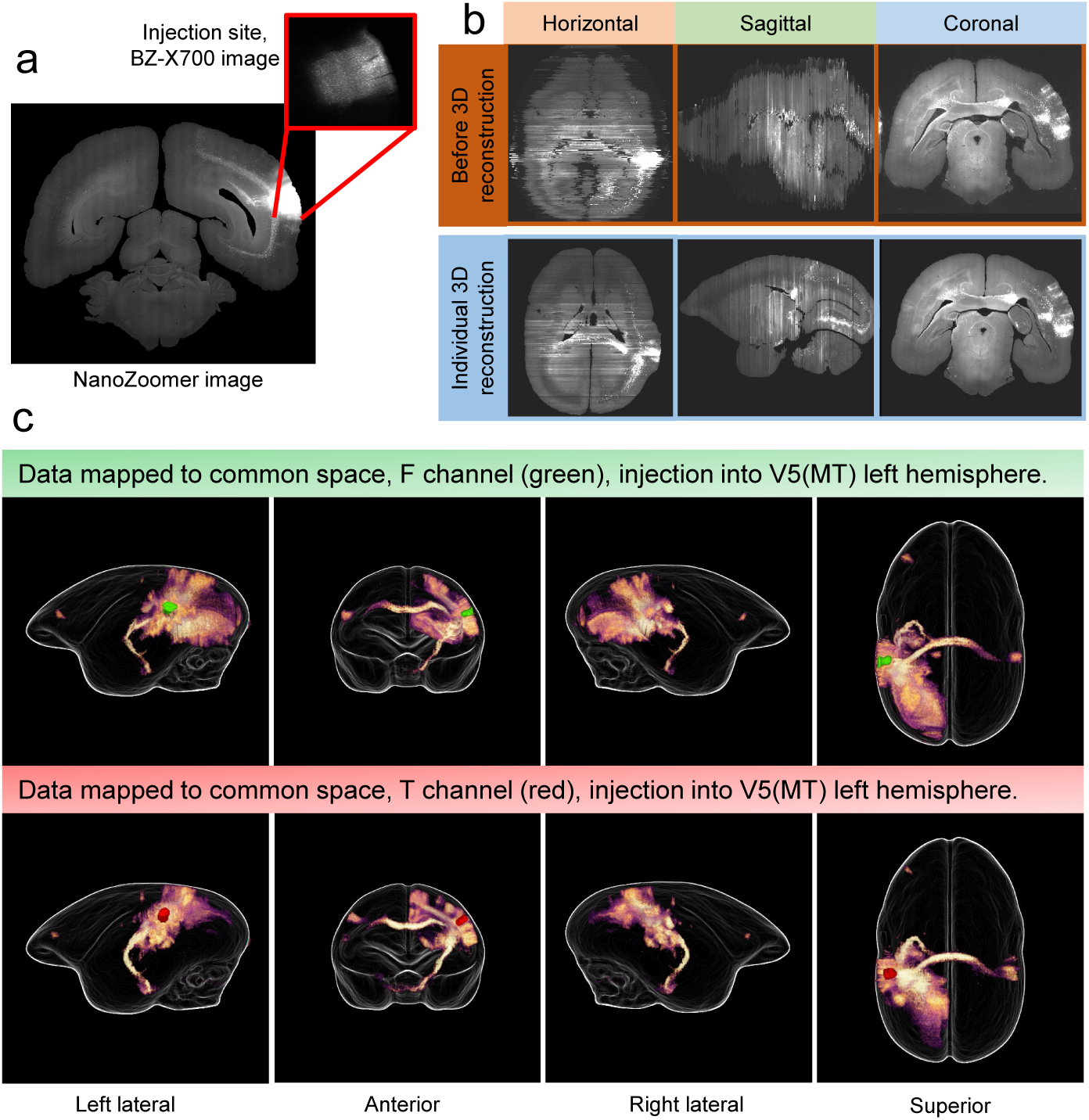
Data from Case 2: (a) An example of a coronal view of the injection site in the NanoZoomer and BZ-X700 images, (b) the 3D reconstruction of an individual brain showing the recovery of the shape by using the block-face reference, (c) the 3D reconstruction of the U-Net segmented fluorescence signal (F and T channels) mapped into the common brain space.

**Fig. 6.**
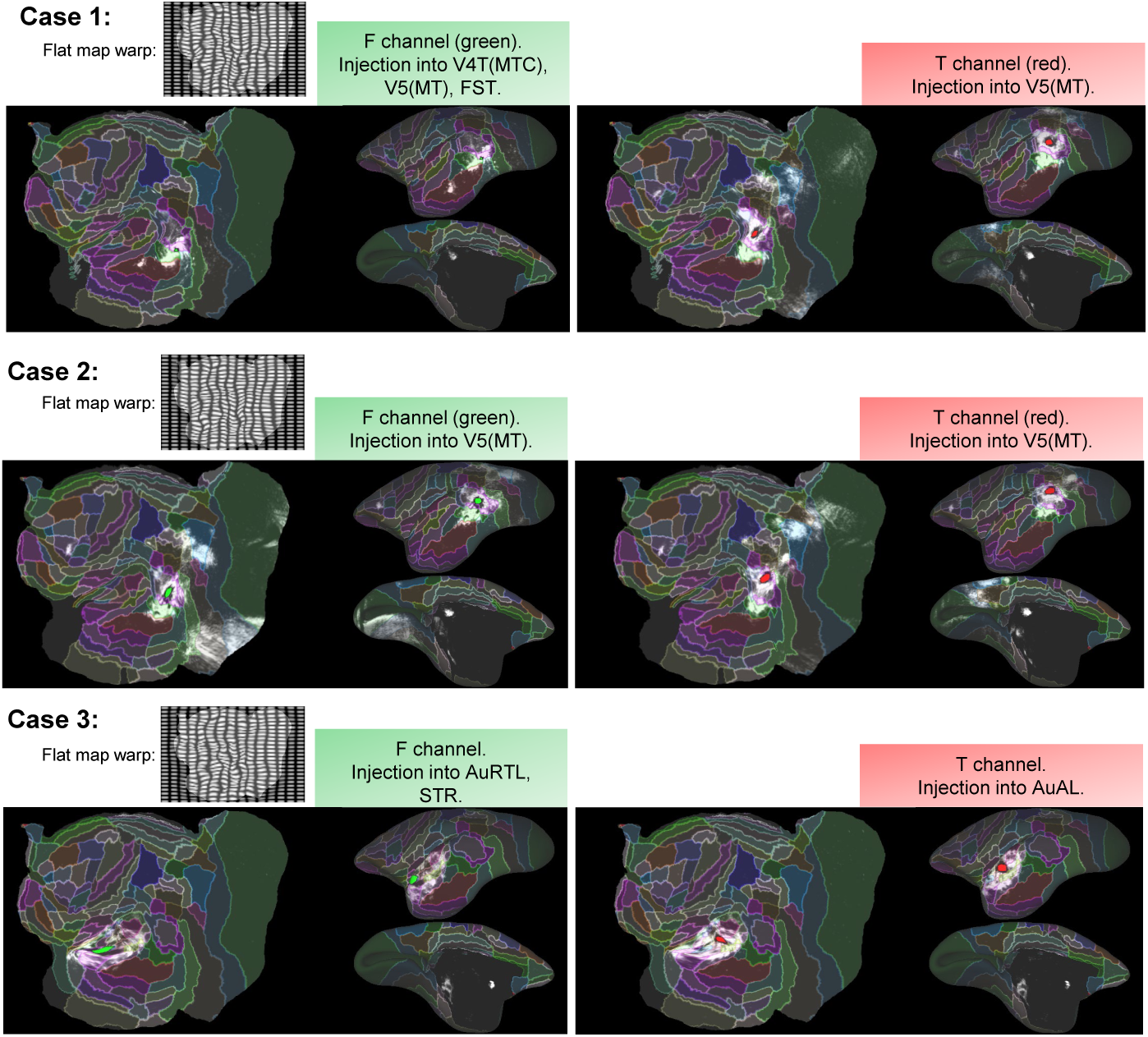
Reconstructions from our pipeline mapped into the common cortical flat map space.

**Fig. 7.**
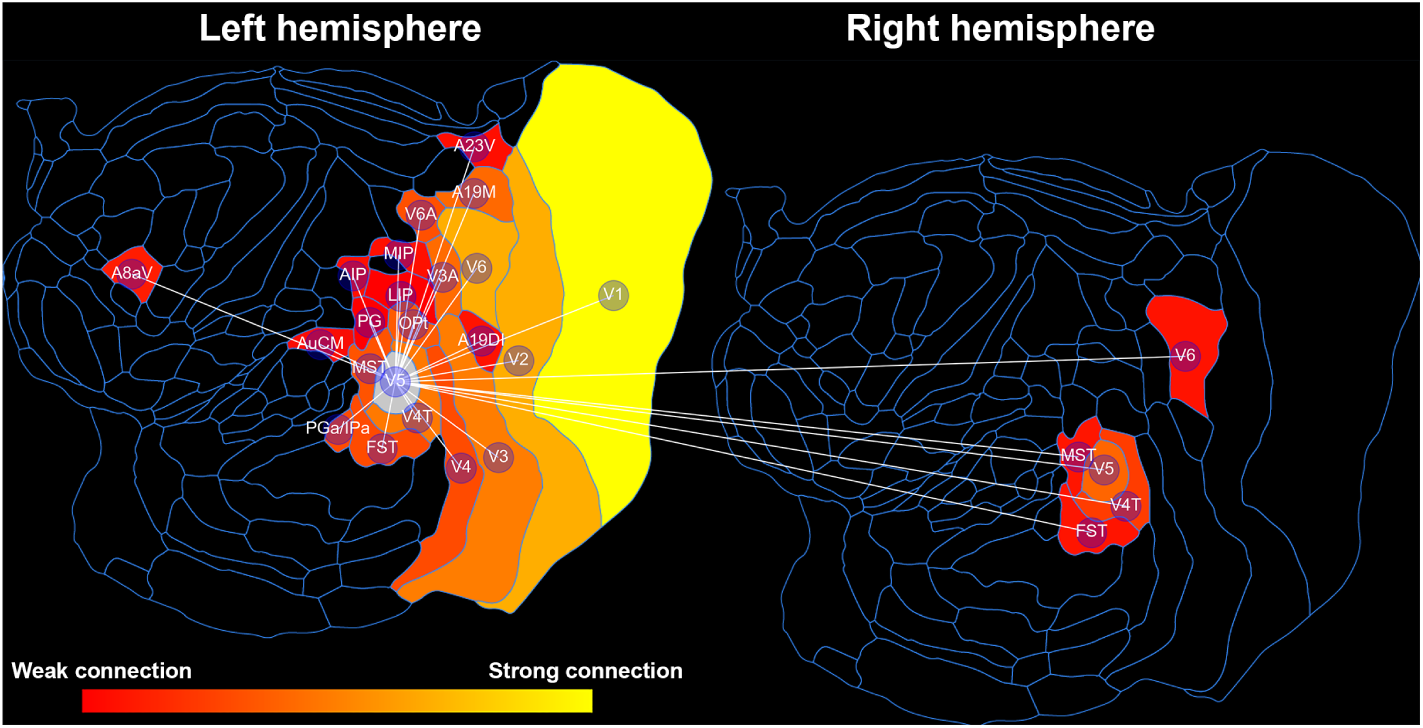
An example connectomics result for Case 2, T channel, injection into region V5(MT) of the left hemisphere. Connectivity strengths between regions were quantified and visualized in the cortical flat map space (showing left and right hemispheres). Values were thresholded to show only strongly connected regions where at least 10% of the region was filled with tracer signal and the color coding was mapped between the maximum and minimum values in the data.

**Fig. 8.**
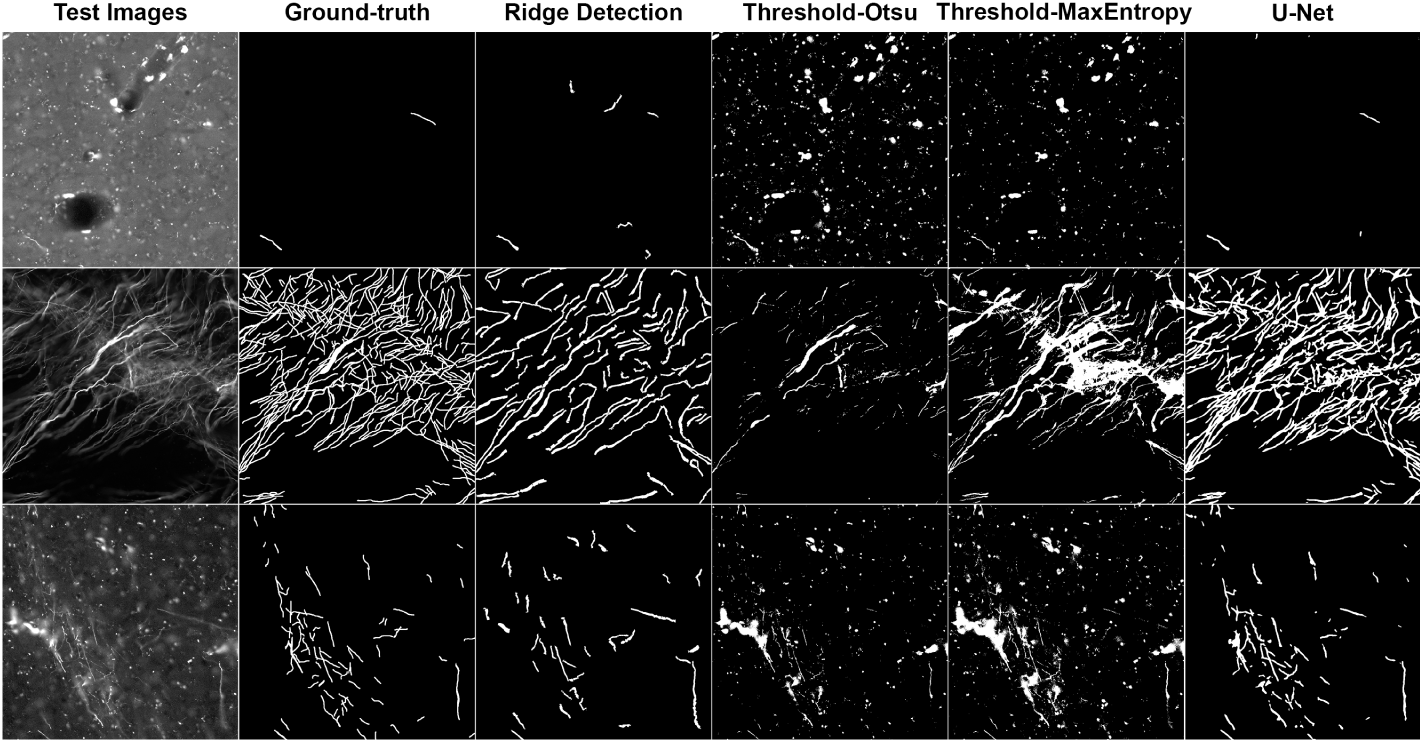
Results of our U-Net tracer segmentation vs other methods (columns) for three images in the test set.

We carried out a quantitative evaluation of the U-Net tracer signal segmentation by comparison with hand drawn ground-truth images and Ridge Detection (Steger (1998)), Otsu thresholding (Otsu (1979)), and Max Entropy thresholding (Kapur et al. (1985)) segmentation methods. Results are shown in Fig. 9, and Table. 1. We computed p-values for our results using the Kruskal-Wallis test (Kruskal and Wallis (1952)) and post-hoc testing was done using Dunn’s test (Dunn (1964)).

**Table 1.** The p-values obtained from Dunn’s test, comparing U-Net and other segmentation methods under all 6 accuracy measures. Gray and light gray represents a p-value that is greater than 0.05 and greater than 0.01 but less than 0.05, respectively.

| Validation method | Dunn's test p-value |  |  |
| --- | --- | --- | --- |
|  | U-Net & Ridge Detection | U-Net & Otsu | U-Net & Max |
| Target overlap | 0.108 | $9.81 \times 10^{-4}$ | $3.97 \times 10^{-3}$ |
| Jaccard index | $4.26 \times 10^{-3}$ | $2.39 \times 10^{-7}$ | $8.68 \times 10^{-7}$ |
| Dice coefficient | $9.36 \times 10^{-3}$ | $6.01 \times 10^{-7}$ | $6.01 \times 10^{-7}$ |
| Area similarity | 0.034 | $4.59 \times 10^{-7}$ | $1.1 \times 10^{-5}$ |
| False negative | 0.108 | $9.81 \times 10^{-4}$ | $3.98 \times 10^{-3}$ |
| False positive | 0.224 | 0.079 | $7.56 \times 10^{-4}$ |

**Fig. 9.**
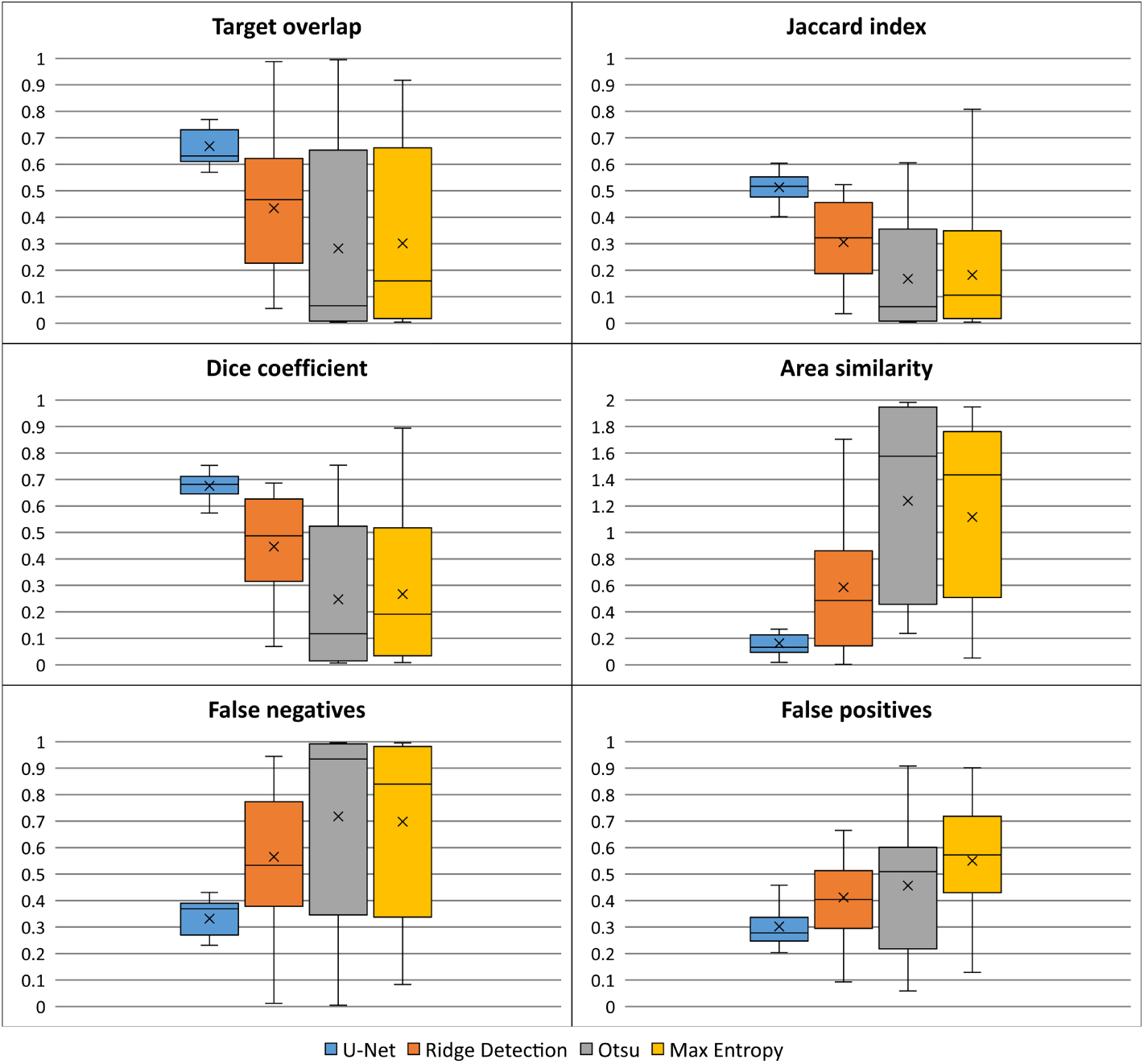
Accuracy of our U-Net tracer segmentation comparing to Ridge Detection, Otsu thresholding, and Max Entropy thresholding, presented using box plots. The top and bottom lines represent the minimum and maximum values, the three middle lines represent the upper quartile, median and lower quartile, respectively and the cross giving the mean. The p-value from the Kruskal-Wallis test is listed below each graph.

We used 20 unseen images of size 1000 × 1000 pixels taken from data from a different brain not used in the U-Net training/validation set. For each of the test images, a ground-truth segmentation image, *G*, was manually drawn. These 20 images were given as input to the U-Net tracer segmentation and other segmentation methods, generating a segmentation image *S*. See Fig. 8 for examples of the results of each algorithm. In each image *r* denotes the region labeled as tracer (binary value 1 with background value 0). Six different measures (taken and adapted from Klein et al. (2009)) were used to assess segmentation accuracy: target overlay=*|S*_*r*_ *∩ G*_*r*_*| / |G*_*r*_*|*, Jaccard index=*|S*_*r*_ *∩ G*_*r*_*|/|S*_*r*_*∪G*_*r*_*|*, Dice coefficient=2*|S*_*r*_*∩G*_*r*_*|/*(*|S*_*r*_*|*+*|G*_*r*_*|*), area similarity=2(*|S*_*r*_*|-|G*_*r*_*|*) */* (*|S*_*r*_*|* + *|G*_*r*_*|*), false negative error=*|G*_*r*_*\S*_*r*_*| / |G*_*r*_*|*, and false positive error=*|S*_*r*_*\G*_*r*_*| / |S*_*r*_*|*, where *G*_*r*_*\S*_*r*_ is the set of elements in *G*_*r*_ but not in *S*_*r*_ etc., and *||* gives the area calculated as a sum of pixels.

The box and whisker plots in all six measures show that our U-Net segmentation outperforms other methods with better accuracy (higher Jaccard index, Dice coefficient, lower false positives and false negatives) and was more consistent (smaller interquartile range with smaller upper and lower bound). Comparatively, the other three methods showed inconsistent performance. For the Jaccard index and Dice coefficient, although the scores were not close to 1 (a perfect match), we still consider the results good due to the ambiguity in drawing the manual ground-truth, where we had to decide on the thickness of the tracer when its boundary was unclear, and decide on how much of the out of focus tracer signal should be considered. Table. 1 points to Ridge Detection performance being close to the U-Net than the other methods (p-value greater than 0.05 in some measures such as false negatives and false positives). But for the two most important segmentation measures, the Jaccard index and Dice coefficient, the U-Net outperformed all other methods at statistical significance (p-value less than 0.01).

The results of 3D registration accuracy are given in Table. 2 where we assessed the accuracy of registration of our 3D reconstructions into the common brain space. We measured the 3D positional difference in corresponding landmarks for the 3D reconstructed fluorescence signal for 10 brains, mapped into the common brain space with the population average MRI contrast. We quantitatively evaluated the registration of fluorescence images since they were the most challenging image type to register, often having weakly visible brain structures in the auto-fluorescence signal to drive image registration. The set of landmarks were based on the definitions in Woodward et al. (2018a). The landmarks and registration error are listed in Table. 2. In the common brain space, where each voxel is 0.1*mm*^3^, the marmoset brain is about 440 voxels in width at the widest point (coronal plane). The average error over all landmarks was calculated to be about 0.39*mm*, or about 4 voxels. Here the average error is in the range of 4*/*440 *≈* 1% of the brain size. This is also reasonable when we consider the difficulty in manually assigning landmarks in the sometimes weak contrast in the brain auto-fluorescence. A previous evaluation of the 2D-to-2D registration of fluorescence to block-face image was carried out in Abe et al. (2017).

**Table 2.**
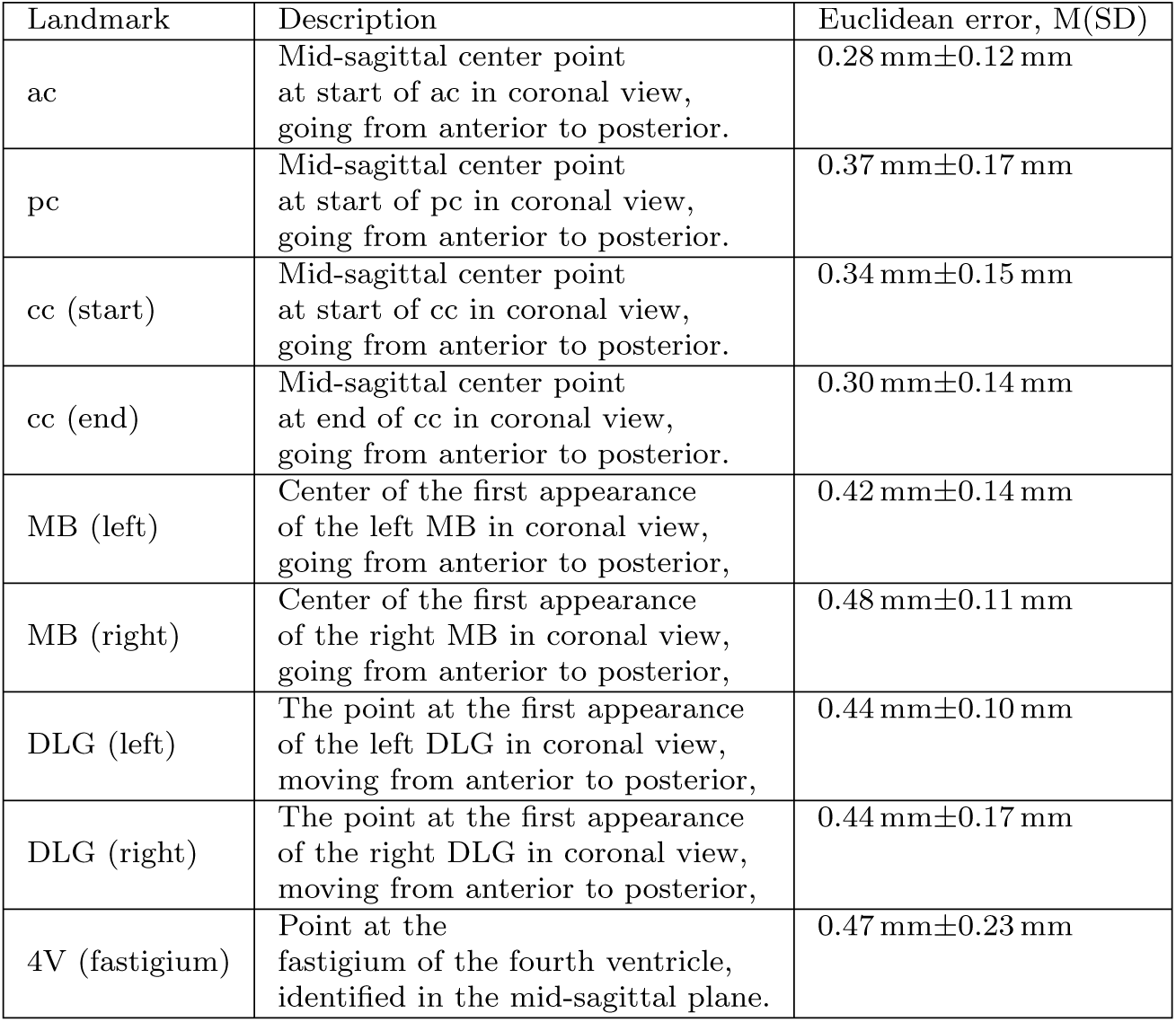
Landmark descriptions and average euclidean error in 3D registration to the common brain space. The anatomical abbreviations are: ac, anterior commissure; pc, posterior commissure; cc, corpus callosum; MB, mammillary body; DLG, dorsal lateral geniculate nucleus; 4V, fastigium of the fourth ventricle. Locations of all landmarks were confirmed visually in horizontal, coronal and sagittal views.

| Landmark | Description | Euclidean error, M(SD) |
| --- | --- | --- |
| ac | Mid-sagittal center point at start of ac in coronal view, going from anterior to posterior. | 0.28 mm±0.12 mm |
| pc | Mid-sagittal center point at start of pc in coronal view, going from anterior to posterior. | 0.37 mm±0.17 mm |
| cc (start) | Mid-sagittal center point at start of cc in coronal view, going from anterior to posterior. | 0.34 mm±0.15 mm |
| cc (end) | Mid-sagittal center point at end of cc in coronal view, going from anterior to posterior. | 0.30 mm±0.14 mm |
| MB (left) | Center of the first appearance of the left MB in coronal view, going from anterior to posterior, | 0.42 mm±0.14 mm |
| MB (right) | Center of the first appearance of the right MB in coronal view, going from anterior to posterior, | 0.48 mm±0.11 mm |
| DLG (left) | The point at the first appearance of the left DLG in coronal view, moving from anterior to posterior, | 0.44 mm±0.10 mm |
| DLG (right) | The point at the first appearance of the right DLG in coronal view, moving from anterior to posterior, | 0.44 mm±0.17 mm |
| 4V (fastigium) | Point at the fastigium of the fourth ventricle, identified in the mid-sagittal plane. | 0.47 mm±0.23 mm |

## 4 Discussion

Our pipeline was able to process different brains concurrently by leveraging an HPC environment. The processing of brain region delineations and entire 3D reconstruction and mapping procedure took about five hours per brain. The main bottleneck was the segmentation of NanoZoomer images using a trained U-Net due to a lack of GPUs. Sequential segmentation of a set of 200 full-resolution fluorescence images for one brain took around a day and a half on a Nvidia RTX 2080 Ti GPU. In total, we can conservatively estimate that we need 2-3 days to process the data for one brain. We consider this reasonable since the experimental data acquisition and work of a neuroanatomist to annotate a brain takes longer than this.

### 4.1 Related work

Comparable state-of-the-art brain mapping activities include: Majka et al. (2016) who describe a method to register marmoset brain data from retrograde tracer experiments into a standard atlas built from Nissl histology slices. In comparison, our experiments used anterograde tracer, we individually annotate each brain, and we have subcortical structures delineated in our common space. We also processed MRI data so that we can make comparisons with diffusion-weighted magnetic resonance imaging (dMRI) based tractography. The retrograde labelled cell detection procedure in their work is manual, while it was necessary for us to develop an automated procedure to segment the dense anterograde tracer signal in our data.

The work of Lin et al. (2019) shares basic similarities with our pipeline by using anterograde tracers and the same NanoZoomer 2.0-HT batch slide scanner to capture fluorescence images. However, they do not use a block-face image as a reference and instead rely on an MRI T2WI of the same brain. In our approach the block-face allows us to directly estimate the true ex-vivo shape of the brain. Relying solely on MRI as a reference means there is more potential for alignment errors since there is no exact correspondence between a fluorescence section and a 2D slice through the MRI volume. Furthermore, they did not use artificial intelligence to segment the tracer signal which may limit the accuracy of connectomics results from their pipeline. Importantly, they used a reference atlas based on a single individual, whereas we overcame this limitation by creating individual brain atlases and mapping them into the common brain space.

For tracer injection studies of other species, the Allen Brain Institute developed a mesoscopic connectome of the mouse brain (Oh et al. (2014)) based on anterograde tracer injections imaged using a TissueCyte machine (Ragan et al. (2012)). In Kuan et al. (2015) they describe the registration procedure, online platform and neuroinformatics tools that were developed to support the production of their connectome. Markov et al. (2014) carried out a tracer injection study for the macaque monkey where results from 29 retrograde tracer injections were analyzed. These have been incorporated into the Cocomac tract-tracing database for the macaque, which uses the results of several hundred tracer study publications (Bakker et al. (2012)). A large number of tracer injection studies have been carried out for the rat brain. Their summary and analysis using network theoretic properties have been investigated in papers such as Swanson et al. (2017) and Bota et al. (2015).

### 4.2 Individual atlasing combined with a common brain space

We took the approach of delineating cortical brain regions for each individual so we could remove the bias of having to use a single template atlas made from a different individual to specify cortical boundaries. This allowed us to precisely map tracer connectivity patterns and ensure the calculation of accurate connectomics. The current limitation of this approach is the manual labor involved in cytoarchitectonically delineating brain boundaries, limiting the scalability of our system. To improve on this we intend to investigate image processing and artificial intelligence techniques for quantitatively defining brain regions from histology. An example of such an approach is the work of Schleicher et al. Schleicher et al. (2005) which extracted and analyzed the change along cortical profiles using the Mahalanobis distance. Even if only a few cortical regions can be identified this can help drive the reconstruction process or be used as a guide to speed up the work of a neuroanatomist. Furthermore, our current cortical delineations can be used as training data for artificial intelligence based cortical region segmentation.

Currently we only drew cortical boundaries and not the sub-cortical structures for each individual. However, the Brain/MINDS atlas defines sub-cortical structures. Through visual inspection we found that the mapping procedure to the common brain space reliably matched the boundaries of sub-cortical regions. This makes it less critical to delineate sub-cortical boundaries for each brain.

A common brain space with atlas has value when we want to integrate data from a number of modalities and when we do not have brain region delineations defined for an individual. Also, having a standard flat map space that we can project data into removes the need to construct an individual flat map for every brain. When histological staining is available, the common atlas can be mapped and used a starting point to refine anatomical boundaries in individual brain data. Additionally, although we can carry out connectomics calculations in the individual brain space, we need a common 3D space for integrating spatial information. For example, anterograde tracer allows us to track the physical path of axonal projections in 3D. Here the common brain space becomes essential for integrating fiber pathways from different tracer experiments.

### 4.3 Tracer signal segmentation accuracy

Results of our U-Net segmentation compared to traditional methods are summarized in Fig. 9 and Table. 1. We were able to successfully segment completely unseen datasets generated from the NanoZoomer and show that artificial intelligence could be used for a challenging image segmentation task. In future work we want to expand our application of artificial intelligence and we have identified a number of targets. First, the background in the block-face and fluorescence images were manually masked and we also segmented the injection side region manually since the number of images to process was minimal. We will attempt to automatically segment out these regions of interest by again using the U-Net approach. The current manually created masks can be used as training data. We also intend to investigate the segmentation of the tracer signal into further features by using multi-class segmentation. For example, if it is possible to characterize synaptic boutons in the tracer signal we can differentiate between passing fiber regions and terminal sites in the data, something that can improve the accuracy of our connectomics calculations.

### 4.4 Accuracy of 3D reconstruction and mapping to common space

Our pipeline can successfully reconstruct the shape of an individual brain for a number of different data types – block-face, tracer fluorescence, injection site, Myelin staining, and individual cortical atlas – and map this data into a common brain space. We can consider the proposed registration procedure to the common brain space to be successful but we will also look at ways to improve this in the future. One approach is to carry out manual adjustments to results in 3D or investigate improved registration using artificial intelligence.

We justify the processing of data at lower image resolution since 2D deformations of fluorescence or Myelin images relative to the block-face occurred from the flattening of the section onto a glass slide. This can be described by low-frequency spatial warping plus a linear offset in position. Since we do not need to process high-resolution images to describe low-frequency spatial deformations we were able to reduce the image size and speed up the computation time for 3D reconstruction and mapping. Since we also saved the transformations at each step we can project any data calculated at lower resolution back into the original high resolution 2D fluorescence images. Also, if more computational resources become available, we can easily increase the resolution at which calculations are done at.

### 4.5 Connectomics

Although we can successfully generate summaries of connectivity across regions – as shown in Fig. 7 – the accuracy needs to be confirmed. Furthermore, a systematic way to treat injections that enter into multiple regions should also be decided upon.

By using anterograde tracer injections we were able to capture the spatial structure of fiber projections, giving us the opportunity to analyze intra-region fiber structure. The U-Net tracer segmentation also allowed us to capture weak tracer signal that might be missed when searching by eye. By mapping the individual atlas back into the original high-resolution fluorescence images we have the opportunity to study the fine-scale tracer projections that might be thresholded out by coarser-grained analyses.

## 5 Conclusion

We have described our connectomics pipeline for the processing and 3D reconstruction of tracer injection images of the common marmoset brain, scanned by a NanoZoomer 2.0-HT batch slide scanner. We developed a number of features that differentiate our work from existing connectomics pipelines. We used artificial intelligence in the form of the U-Net deep network architecture to precisely segment the tracer signal from the background, even when images were noisy and contained bright background features. We also cytoarchitectonically delineated the cortical regions of each brain to create accurate 3D individual atlases. This allowed us to avoid bias from using a template brain atlas to specify cortical boundaries. We created a common brain space based on a population average ex-vivo MRI T2WI contrast mapped with the Brain/MINDS marmoset atlas and mapped individual data into it. To further normalize our data we projected it into a common 2D cortical flat map space and precisely aligned individual cortical regions with those defined in the common brain space.

Now that we have established a connectomics pipeline we intend to process a number of brains and calculate a connectomics matrix of anatomical connectivity for the injected brain regions. We intend to integrate our connectomics data with those from other tracer study pipelines (both anterograde and retrograde) to get closer to a complete understanding of the structural connectivity of the common marmoset brain. In the future we also plan to use our pipeline to develop a population-based brain atlas where we will incorporate histological annotation and information derived from structural MRI and DWI data. This will allow us to evaluate the variation of cortical region boundaries across individuals.

## Acknowledgements

We thank Akiya Watakabe for sharing his expertise in marmoset tracer injection studies; Masahide Maeda for his technical support in maintaining the server used in this work; and Carlos Enrique Guiterrez and Hiromichi Tsukada for their participation and contribution in previous tracer study activities.

## Compliance with ethical standards

### Research involving human participants and/or animals

All experimental procedures were approved by the Experimental Animal Committee of RIKEN, or by the Experimental Animal Committee of the National Center of Neurology and Psychiatry. The marmosets were handled in accordance with the Guiding Principles of the Care and Use of Animals in the Field of Physiological Science formulated by the Japanese Physiological Society.

### Funding

This research was supported by the program for Brain Mapping by Integrated Neurotechnologies for Disease Studies (Brain/MINDS) from the Japan Agency for Medical Research and Development, AMED, grants JP19dm0207001 and JP19dm0207088.

